# Pear flower and leaf microbiome dynamics during the naturally occurring spread of *Erwinia amylovora*

**DOI:** 10.1101/2024.01.08.574646

**Authors:** Aia Oz, Orly Mairesse, Shira Raikin, Hila Hanani, Mery Dafny Yelin, Itai Sharon

**Author notes:** These authors contributed equally to this work (Aia Oz, Orly Mairesse).

## Abstract

*Erwinia amylovora* is the causal pathogen of fire blight, a contagious disease that affects apple and pear trees and other members of the family Rosaceae. In this study, we investigated the population dynamics of the pear flower microbiome in an agricultural setting during the naturally occurring infection of *E. amylovora*. Five potential factors were considered: collection date, the flower’s phenological stage, location on the tree, location within the orchard, and pear cultivar. The phenological stage and the collection date were identified as the most important factors associated with pear flower microbiome composition. The location of the tree in the orchard and the flower’s location on the tree had a marginal effect on the microbiome composition. The leaf microbiome reflected that of the abundant phenological stage on each date. The flower microbiome shifted towards *E. amylovora,* dominating the community as time and phenological stages progressed, leading to decreased community diversity. The strain population of *E. amylovora* remained similar throughout the entire collection period. In contrast, other taxa, including Pseudomonas, Pantoea, Lactobacillus, and Sphingomonas, were represented by dozens of amplicon sequence variants (ASVs), and different succession patterns in their populations were observed. Some of the taxa identified include known antagonists to *E. amylovora*. Overall, our results suggest that flower physiology and the interaction with the environment are strongly associated with the pear flower microbiome and should be considered separately. Strain succession patterns for the different taxa under *E. amylovora* spread may help in choosing candidates for antagonist-based treatments for fire blight.

**Importance:** The spread of pathogens in plants is an important ecological phenomenon and has a significant economic impact on agriculture. Flowers serve as the entry point for *E. amylovora,* but members of the flower microbiome can inhibit or slow down the proliferation and penetration of the pathogen. Knowledge about leaf and flower microbiome response to the naturally occurring spread of *E. amylovora* is still lacking. The current study is the first to describe the flower microbiome dynamics during the naturally occurring infection *of E. amylovora*. Unlike previous studies, our experiment design enabled us to evaluate the contribution of five important environmental parameters to the community composition. We identified different strain succession patterns across different taxa in the flower consortia throughout the season. These results contribute to our understanding of plant microbial ecology during pathogen spread and can help to improve biological treatments for these diseases.

## Introduction

Microbes inhabit nearly all parts of the plant, from roots to leaves and flowers, both on the plant surface and within it. Microbes that live on the root surface (the rhizoplane microbiome), inside the roots (endophytes), and in the soil near the roots (the rhizosphere microbiome) contribute to the plant’s health and growth (1–3). The phyllosphere microbiome includes microbes that live on leaves, flowers, fruits, and seeds. Microbes inhabiting the phyllosphere contribute to plant health, positively interact with pollinators, and promote plant growth (1,4,5), but can also cause several plant diseases (6).

Flowers serve as a bacterial entry point to the plant. Their high nutrient content attracts pollinators that serve as vectors (7) for microbial colonization and succession (8,9). The flower microbiome can impact the nectar composition (10) and is suspected to play a role in attracting pollinators (11). The phyllosphere microbial composition is affected by plant species, genetics, age, and various environmental conditions such as agricultural management, weather conditions, and soil chemistry (12). The flower microbiome differs even for closely related species such as pear (*Pyrus communis*) and apple (*Malus domestica*). This may be due to the different flower and nectar characteristics, such as pH and sugar content (13). Despite decades of research on flower-associated microbes, the interactions between the plant and the phyllosphere microbiome (12) and the drivers that most affect the flower microbiome dynamics still need to be fully understood (8).

The pathogenic, gram-negative enterobacterium *Erwinia amylovora* uses the floral nectar secretory cells as an entry point when infecting rosaceous trees in fire blight disease (14,15). Fire blight is a necrotic plant disease affecting apple and pear orchards worldwide, causing severe economic damage (16,17). Infection results in blights of the shoots, fruit, and rootstock, lowering crop yields and sometimes killing entire orchards (14). Following infection, *E. amylovora* moves through the vascular tissue to colonize the bark and the woody parts of the tree, thus causing blight and creating a bacterial reservoir for future infections (15,17,18). Several environmental and physiological conditions are known to affect the spread of *E. amylovora* in orchards. The movement of the bacteria in the shoots, branches and barks is influenced by tree age, growth rate, pruning methods and tree design (19). The growth rate is affected by the amount of fertilizer applied, and treatments with plant growth regulators. The initial *E. amylovora* reservoir in the orchard and its vicinity is affected by infections in the previous years and the sanitation quality to remove infected branches. Finally, weather conditions that favor *E. amylovora* are critical to the disease infection. These mainly include warm temperatures and wetness in the form of rain or dew during the bloom period (14,15,20–23). Current knowledge on microbiome dynamics during the spread of *E. amylovora* in Rosaceous orchards is based on inoculation experiments in which apple flowers were sprayed with *E. amylovora*. These experiments revealed that members of the families Enterobacteriaceae and Pseudomonadaceae dominate the non-infected flowers, with only small amounts of *E. amylovora* detected. In contrast, the infected flowers were dominated by the inoculated strain (9,24). Preventive treatments for fire blight are based on antibiotics, copper compounds, and antagonistic bacteria, all with environmental impacts and limited success (17,25–28). Understanding the population dynamics and the factors affecting them might help to develop and improve effective, non-destructive methods for treating fire blight.

Here, we aim to explore the effect of different factors on the pear flower microbiome during *E. amylovora* natural infection. We evaluated the contribution of the phenological stage, collection date, location in the orchard, and location on the tree on the microbiome of pear flowers and leaves using 16S small subunit (SSU) rRNA amplicon surveys. Coincidentally, the orchard has undergone an *E. amylovora* infection, which developed into a fire blight event observed later in the season. To our knowledge, no study described flower microbiome dynamics in untreated pear orchards following the natural occurrence of *E. amylovora* infection. Our study documents, for the first time, the factors that affect the natural dynamics of the Rosaceous flower microbiome during *E. amylovora* infection.

## Results

### Experiment setup

We aimed to study the factors influencing pear flower microbiome dynamics during an *E. amylovora* infection. To this end, flower and leaf samples were collected during the spring of 2019 from an orchard in the Hula Valley, northern Israel. The study site was selected because it is at high risk for fire blight due to the climatic conditions in the region. Several events of naturally occurring fire blight were documented in the region in the years preceding our study. Five potential determinants were considered: collection date (time), phenological stage, location in the orchard, location on the tree, and cultivar. To investigate the impact of the first four variables, we designed a collection strategy that includes collecting flower and leaf samples from six pear (*Pyrus communis*) Coscia trees on five dates. To assess the impact of the phenological stage on microbial composition, we considered the following five phenological stages: white buds; initial bloom, characterized by open flowers with pink, non-mature anthers; full bloom, in which the anthers matured and opened, and changed color to black; initial petal fall; and complete petal fall (**Fig. 1A**). For each collection date, two to four abundant phenological stages were selected for sampling. The flower samples were collected from two groups of trees located ∼15m apart, at two heights on each tree (high=∼2.5m, low=∼1.5m), resulting in a total of 12 samples per phenological stage per collection date (**Fig. 1B** and Methods). Young leaf samples were collected from each tree on each date, totaling six leaf samples per collection date. To investigate the effect of the cultivar, we collected flowers and leaves from six Spadona trees on three dates earlier in the season, one phenological stage on each date. The Spadona samples were used to compare the microbiome of the white buds and initial bloom stages in the two different cultivars. The samples underwent 16S amplicon sequencing (V3-V4 region; see Methods), with an average sequencing depth of ∼33K paired reads per sample. We identified six abundant Buchnera amplicon sequence variants (ASVs) that cover >97% of the Buchnera populations, in Spadona flowers only. These ASVs align at 100% similarity to the 16S gene of the endosymbiont *Buchnera aphidicola*. This suggests that aphids (or their eggs) were present on the flowers even though we did not observe aphids during the collection period. To our knowledge, *Buchnera* is not a part of the flower microbiome, and we, therefore, removed it from our analyses. Overall, a total of 223 samples (176 flowers and 47 leaves) were collected and sequenced (**Table S1**). The samples were rarefied separately for each experiment (see **Table S2**).

**Figure 1:**
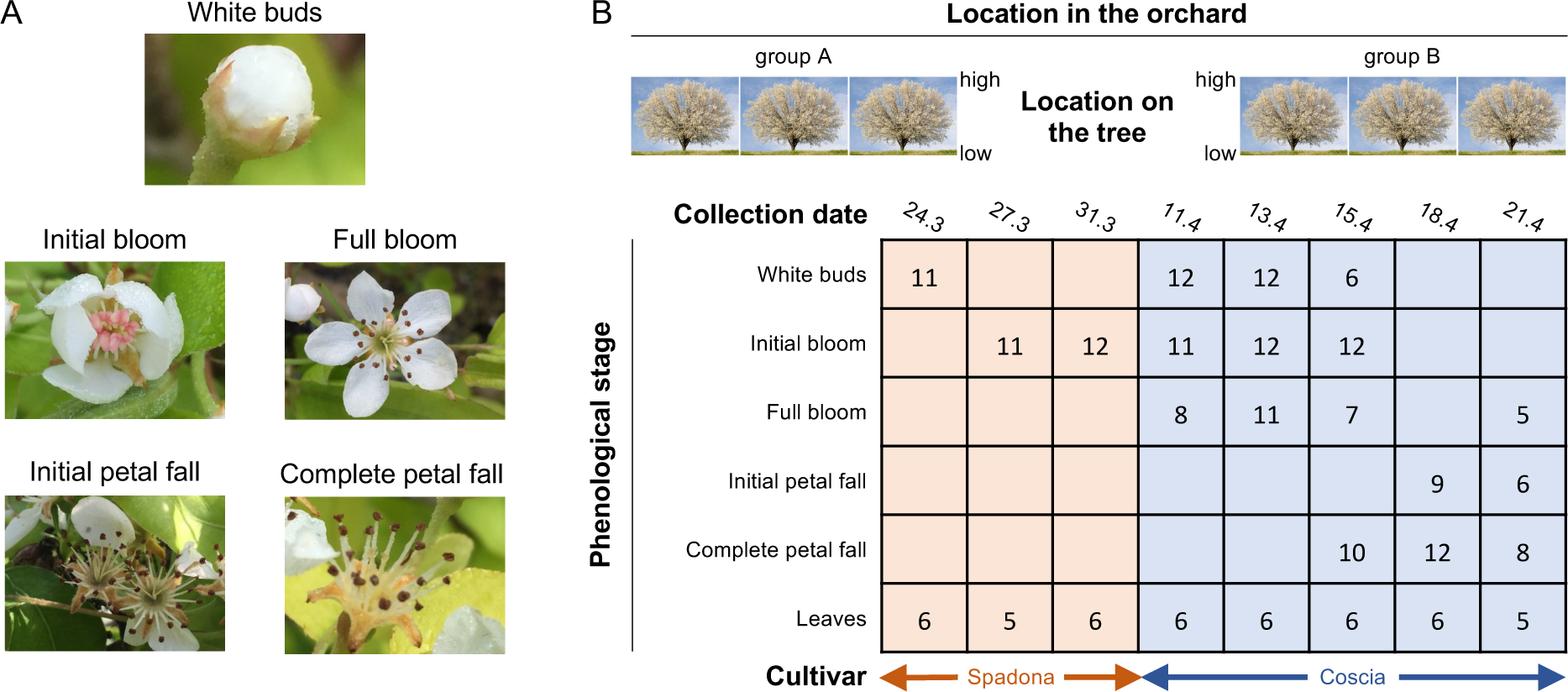
a summary of the collection strategy. (A) The flower phenological stages considered in this study. (B) Five variables were considered: location in the orchard, location on the tree (high or low), collection date, phenological stage and cultivar. The numbers indicate the number of samples collected at each collection date/phenological stage. The number of samples used in different analyses may vary according to the rarefaction threshold used.

### The effect of the five variables on the microbiome

A total of 29 different phyla were identified in the flower and leaf samples of the Spadona and Coscia cultivars (**Table S3, Fig. S2**). Proteobacteria is the most abundant phylum in both the Spadona and Coscia samples, with an average abundance ranging from 60.7% of the population (11.4, Coscia, initial bloom) to 99.5% (21.4, Coscia, complete petal fall). Other abundant phyla include Firmicutes (now Bacillota), Bacteroidota, and Actinobacteriota. These phyla were previously reported to be the most abundant in pear flowers, although in different proportions (13). Within the Proteobacteria, a significant shift between represented genera is observed when comparing earlier dates and phenological stages (Pseudomonas, Lactobacillus, Sphingomonas, Methylobacterium-Methylorubrum) to later ones, which are dominated by Erwinia (see later). These findings are in line with previous studies of Rosaceous trees, in which the families Enterobacteriaceae and Pseudomonadaceae (phylum Pseudomonadota, previously Gammaproteobacteria), Bacillaceae and Lactobacillaceae (phylum Bacillota, previously Firmicutes), and the phylum Candidatus Saccharibacteria (previously TM7) were reported (8,9,24,29).

In our study site, both groups of trees were exposed to similar environmental conditions, such as sun exposure. We, therefore, anticipate that factors that affect the microbiome at different heights on the tree will have a similar effect on microbiome diversity at different locations in the orchard. Also, we assume that local factors that can affect the microbiome at different locations in the orchard, such as wind and pollinators, will similarly affect flowers and leaves at different heights on the tree. To this end, we compared microbiome diversity at different heights while ignoring the location in the orchard and vice versa.

The flower samples collected from different heights in Coscia trees show negligible and marginally significant partitioning when confounding for both the collection date and the phenological stage of the flowers (R^2^=0.01, p-value=0.05; PERMANOVA, **Table S4**). The Shannon and Chao1 diversity indexes at different heights are statistically insignificantly different when comparing different heights over the same phenological stages and dates (Kruskal-Wallis, **Fig. 2A**). The location in the orchard had a small yet significant effect on both the flower and leaf microbiome when confounding for both collection date and phenological stage (R^2^=0.01, p-value=0.002; PERMANOVA, **Table S4**). Differences in alpha diversity for the location in the orchard were not statistically significant (Kruskal-Wallis, **Fig. 2B**). In Spadona, the location in the orchard showed a higher yet marginally significant effect on the microbiome (R^2^=0.1, p-value=0.03; PERMANOVA, **Table S4**). A Constrained canonical analysis of principal coordinates (CAP) of the Coscia flowers shows that the contribution of the height and location in the orchard to the first two dimensions is small, supporting the conclusion that the effect of these variables is small (**Fig. 3A**). Therefore, we disregarded these variables in the other analyses.

**Figure 2:**
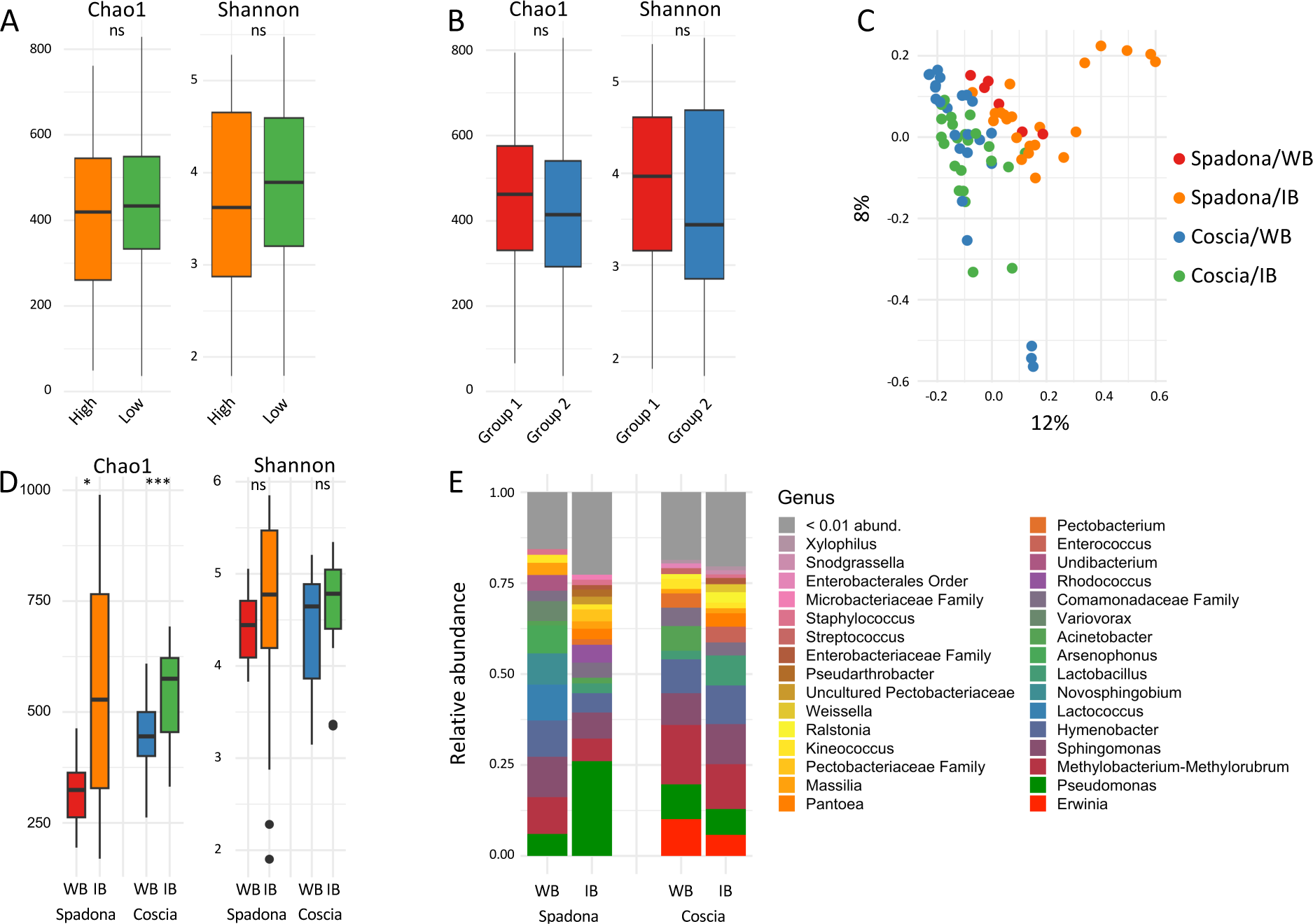
(A) Distribution of Chao1 and Shannon indexes for samples collected at different heights, and (B) The same, for samples collected at different locations in the orchard. Differences between all pairs are insignificant (Kruskal-Wallis). (C) PcoA plot of the white buds (WB) and initial bloom (IB) samples from Spadona (24.3, 27.3, 31.3) and Coscia (11.4 and 13.4). (D) Chao1 (left) and Shannon (right) diversity indexes of the samples, grouped by cultivar and phenological stage. Differences in Shannon diversity between the phenological stages and the cultivars are insignificant (Kruskal-Wallis). The Chao1 index is significantly different (p-value=0.0025, Kruskal-Wallis). (E) Relative abundances of genera in the white buds and initial bloom phenological stages, Spadona and Coscia.

**Figure 3:**
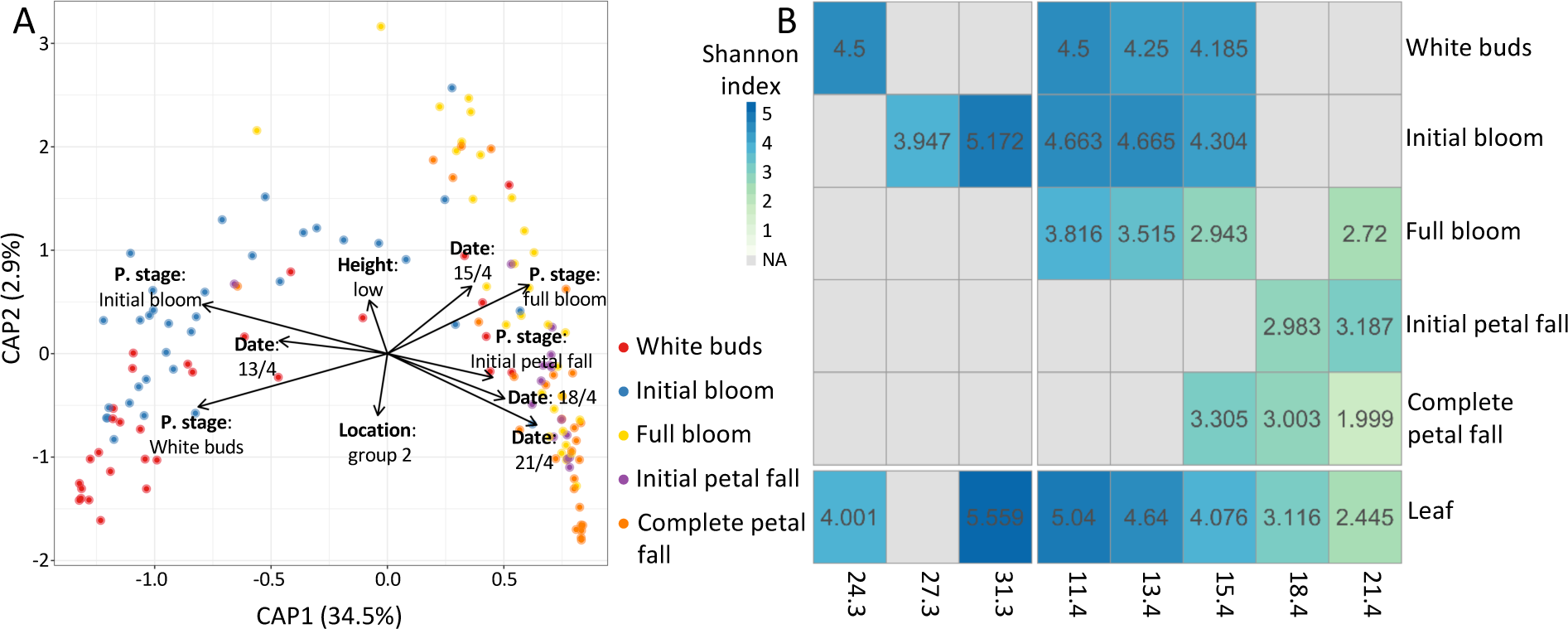
(A) Constrained canonical analysis of principal coordinates of Coscia flower samples, using Bray-Curtis dissimilarity. The model applied was: phenological stage + collection date + height + location in the orchard. The plot is colored by the phenological stage. (B) Mean Shannon diversity over the same dates and phenological stages.

We compared the samples collected from the Spadona cultivar to those from the same phenological stages in Coscia. In this analysis, only samples collected during the first two collection dates of Coscia (11.4, 13.4) were analyzed due to the spread of *E. amylovora* in the orchard at later dates (see below). Principal coordinates analysis (PCoA) shows that samples from the two cultivars cluster separately while samples of different phenological stages from the same cultivar cluster together (**Fig. 2C**). In both cultivars, community diversity increased from the white buds to the initial bloom stages (Chao1, not Shannon, **Fig. 2D**). Notable differentially abundant genera between Spadona and Coscia include the genera Erwinia, Lactococcus, Rhodococcus, and Pseudomonas (**Fig. 2E**). We note that since the samples were collected on different dates, it is impossible to determine whether the observed differences are because of the cultivar or the environmental conditions.

We considered the effect of the phenological stage and collection date through a PERMANOVA model that includes both variables and the interaction term. The collection date (R^2^=0.25, p-value=0.001), the phenological stage (R^2^=0.17, p-value=0.001) and the interaction term (R^2^=0.06, p-value=0.003) all have a significant effect on the microbiome. Community composition demonstrated a shift from the first two to the last three phenological stages (**Fig. 3A**, **S3**A), and from the first two to the last two collection dates with the middle collection date (15.4) representing a transition between the two groups (**Fig. 3A**, **S3**B). The large effect of the phenological stage and the collection date on the microbiome compared to the location on the tree and in the orchard is also demonstrated by the CAP of the Coscia flowers (**Fig, 3**A). Community diversity reached its peak during the first two phenological stages and declined sharply with the spread of *E. amylovora* (**Fig. 3B**). Community diversity was significantly different for samples that were collected in different dates for the full bloom and complete petal fall phenological stages, and for leaves, but showed no significant differences for the other phenological stages. Community diversity was significantly different for most samples of different phenological stages that were collected in the same dates (all dates except 18.4, with only two phenological stages collected) (**Table S5** and **S6**). The leaf microbiome diversity changed significantly over time and was overall similar to the one of the dominant phenological stage at each date (**Fig 3B**). These differences are mostly due to the spread of *E. amylovora* (discussed later). A breakdown of the samples to phenological stages at each date, and to collection dates at each phenological stage, shows distinct clustering of groups that are dominated by *E. amylovora* and those that are not (**Fig. S4**, **S5** and **S6**). These groups also differ in community diversity (**Fig. 3B**, **4A**), demonstrating the influence of both phenological stage and collection date on the microbiome composition.

### The spread of *Erwinia amylovora* in the orchard

The orchard sampled has a history of fire blight caused by natural infections and inoculation experiments performed in previous years. During the collection season, optimal conditions for the spread and proliferation of *E. amylovora* occurred several times (**Fig. S1**). With the progression of the date and the phenological stage, the flower and leaf microbial communities became dominated by members of the family Erwiniaceae.

Assigning taxonomy to community members based on 16S amplicon sequencing is limited by the resolution of the taxonomic classifier used. The SILVA classifier that we used classified the Erwiniaceae at the family level. To evaluate the large Erwiniaceae populations in our samples, we further analyzed all Erwiniaceae ASVs in our data by comparing them to the nt database of NCBI. We identified eight ASVs that dominated the Erwiniaceae population, with six aligning at 100% and two aligning at 99.35% and 99.75% to the 16S gene of *E. amylovora*. The second best hit with complete taxonomy other than *E. amylovora* was usually *Erwinia pyrifoliae*, another blight-causing pathogen in pears (30). Other Erwiniaceae hits aligned at <99% similarity in all cases. We therefore concluded that these eight ASVs represent *E. amylovora* strain(s) in our study. An additional Erwiniaceae ASV aligned at 96.36% to *E. amylovora* and was not considered a representative of the species.

In Coscia flowers, the relative abundance of *E. amylovora* ASVs increased significantly with the progression of both the collection date and the phenological stage until reaching ∼95% of the community in the last date and phenological stage. The relative abundance of *E. amylovora* on leaves increased steadily to >85% in the last collection date (**Fig. 4A**).

**Figure 4:**
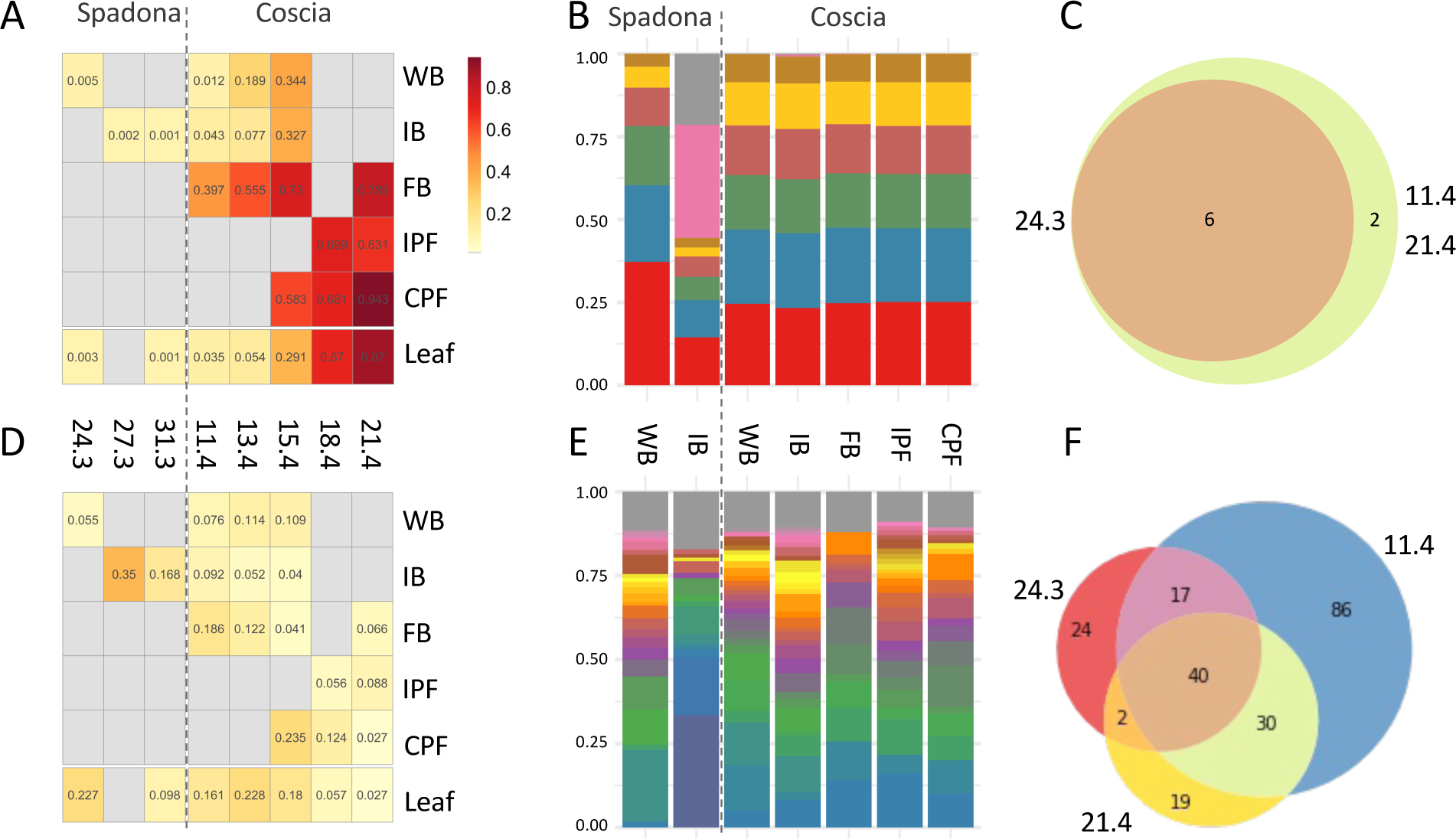
(A) mean relative abundances of ASVs assigned to *E. amylovora* in flowers and leaves across the different collection dates and phenological stages. WB=white buds, IB=initial bloom, FB=full bloom, IPF=initial petal fall, CPF=complete petal fall. (B) The distribution of *E. amylovora* ASVs in each phenological stage. (C) The number of Erwiniaceae ASVs shared between samples collected on the first collection date for Spadona (24.3), the first collection date for Coscia (11.4), and the last collection date (21.4). (D-F) Same as A-C for Pseudomonas.

The six ASVs aligned with the 16S gene of *E. amylovora* at 100% similarity were identified in all collection dates of the Spadona and Coscia flowers (**Fig. 4B** and C). The genome of *E. amylovora* includes seven copies of the 16S gene that differ by a few nucleotides; therefore, it is impossible to determine whether the ASVs represent one or multiple strains. Other lineages, including Pseudomonas, Pantoea, Lactobacillus, and Sphingomonas, declined in their relative abundance with the spread of *E. amylovora* (**Fig. 4D** and **S4**). These taxa were represented by dozens of ASVs, with certain ASVs present only in specific time points and others detected in the first and last collection dates and throughout the collection period (**Fig. 4B** and C, **Fig. S7,** and **Fig. S8**). These trends suggest strain succession had occurred during the collection period, possibly as a response to the spread of *E. amylovora*. The alpha diversity of Pseudomonas populations in each sample did not change significantly throughout the collection period despite their decline in relative abundance. On the other hand, the diversity of Sphingomonas declined, and the diversity of Lactobacillus demonstrated mixed trends (**Fig. S7** and **Fig. S8**).

## Discussion

The flower microbiome has important roles in the plant life cycle. Multiple factors may affect its composition and activity, and understanding these factors can help improve prediction and treatment strategies and develop antagonist-based treatments against diseases such as fire blight. Studying the flower microbiome is challenging due to a multitude of technical reasons: separating the microbial DNA from the DNA of the host plant is difficult, and mitochondrial and chloroplast 16S fragments are amplified by most universal primers in amplicon sequencing. Also, in most cases, the flowering season is short, and experiments cannot be repeated in the same year. In this study, we describe the pear flower microbiome during the spread of *E. amylovora*. The weather conditions in the orchard during the study period favored fire blight disease, and disease signs were observed late in the flowering period. To our knowledge, only a few amplicon-based studies targeted the pear flower microbiome, with most studies on the Rosaceous flower microbiome focusing on apple trees. The effect of the *E. amylovora* population on the leaf and flower microbiome and the dynamics of the plant microbiome during the spread of *E. amylovora* were mostly studied through culturing and inoculation experiments. To our knowledge, the current study is the first to describe microbial community dynamics during a naturally occurring spread of *E. Amylovora*.

We aimed to reveal the contribution of five factors to the pear flower microbiome dynamics: the height of the flowers on the tree, the location of the tree in the orchard, the phenological stage, the collection date, and the cultivar. Our results demonstrate small yet significant differences in the flower microbiome between different heights despite a relatively small difference in the heights compared. These differences may be due to continuous differences in the flowers’ exposure to environmental conditions such as sun hours and wind that can directly affect temperature and humidity. The turn of the tree, which was not tested in this study, may also receive different environmental conditions that can reduce wetness in the morning hours and affect the flower microbiome. The microbiome of trees at different locations in the orchard was overall similar but exhibited small yet significant differences. We attribute these differences to local environmental factors and microbial community processes that affect different parts of the orchard and are noticeable even over distances of 10-20 meters. Pollinators, which were observed and active in the orchard during the collection period, are known contributors to microbial dynamics (7). Their preferences and behavior may also contribute to differences across locations in the orchard and height, but these were not considered in the current study.

Farmers often use plant growth regulators in commercial orchards to make Spadona and Coscia cultivars bloom simultaneously to get optimal pollination and high yield. However, in our experiment plot, minimal chemical intervention was applied which resulted in different flowering periods for the two cultivars. Our results show that Spadona and Coscia cultivars have slightly different microbiomes during the first two phenological stages (white buds and initial bloom), which were tested in this study. The Spadona trees bloomed 2-3 weeks before the Coscia trees during the 2019 flowering season studied here. Therefore, the differences are probably due to different environmental conditions. In apples, the cultivar did not significantly affect the microbial composition when compared during the same period (29).

The phenological stage and the collection date are interrelated but represent different factors. The phenological stage represents the flower physiology (*e.g*., the exposure of the flower stigma), nectar availability and composition, and flower age. The collection date is related to external factors, in particular, weather and exposure to pollinators. Our experiment was heavily influenced by the spread of *E. amylovora* in the orchard, which became the most abundant bacterial species in the flower and leaf communities. Still, flowers collected at different phenological stages over the same dates exhibited distinct community structures, as did flowers in the same phenological stage collected over different dates. These differences were significant and highlight the need to consider both factors when designing experiments aimed at studying the flower microbiome. Interestingly, the results show that leaf microbial populations reflect the dominant phenological stages in each date. Leaf age, which was not considered in our experiments, may play a role in determining the leaf microbial composition, as other environmental conditions, such as the weather.

In most phenological stages and cultivars, >99% of the community was dominated by four phyla: Proteobacteria, Firmicutes (now Bacillota), Bacteroidota, and Actinobacteriota. These phyla were also reported previously in studies of apple and pear flowers (13). Proteobacteria account for 60-99% of the community in all date and phenological stage combinations. Proteobacteria was previously reported to account for a similar proportion of the community in apple flowers (9,24,29) and wildflowers (31), with mostly similar abundant families to those reported in our study.

The flowering season in the late spring has climatic conditions favored by *E. amylovora*. The optimal temperature for *E. amylovora* proliferation is 28ºC, despite being able to survive and multiply in low temperatures (32). The proliferation of *E. amylovora* also depends on water availability provided by rain or dew (15). The 2019 flowering season had 14 days (out of 45) that met the criteria of the "Fire Blight Control Advisory" decision support system for fire blight high-risk weather (33). The pathogen, which was present at ∼0.5% of the population in the Spadona samples, occupied close to 95% of the population in the last collection date and phenological stage. Fire blight symptoms were not observed in the orchard during the flowering period but rather, as expected, only a few weeks later. The prevalence of *E. amylovora* differed across phenological stages and collection dates, in line with previous studies (20,33). Community diversity increased from the white buds to the initial bloom stages before the spread of *E. amylovora* but then declined until the last phenological stage and collection date following the spread of *E. amylovora*. These dynamics are expected and are different from those observed in healthy bacterial communities of apples and pears (8,13). The same six *E. amylovora* ASVs were observed in similar proportions throughout the entire collection period, suggesting that the infecting strain(s) were present in the orchard before the flowering period and that no strain turnover occurred during the flowering season. In contrast, other abundant taxa, such as Pseudomonas, were represented by a larger pool of ASVs, with some present over the entire period and others detected at certain time points only.

In contrast to the microbiome diversity, the dynamics of population diversity for the different taxa varied with some groups, such as Pseudomonas, maintaining a similar diversity even when the group’s share in the population decreased, while the diversity of others, such as Sphingomonas, declining with time progression. Previous studies showed that Pseudomonadaceae was negatively correlated with *E. amylovora* when the microbiome of single flowers was tested (9,24). In our study, samples consist of 30 flowers, thus representing an average over multiple blossoms. Therefore, it is not possible to determine whether *E. amylovora* and the other taxa are evenly spread over the different flowers, or if the communities of specific flowers greatly differ.

## Conclusion

In our experiment, the spread of *E. amylovora* in the orchard occurred from one or a few strains that were present in the orchard from the beginning of the flowering season. No turnover in the *E. amylovora* population has been observed. At the same time, the share of the pathogen in the community increased from less than 1% of the flower microbiome on the first collection date to more than 94% on the last date and phenological stage. Other groups, such as Pseudomonas, Sphingomonas, and Lactobacillus, declined in overall abundance but were represented by dozens of strains with significant turnover throughout the season. The collection date and the phenological stage both explain a significant portion of the microbiome dynamics during the spread of *E. amylovora*, while the cultivar, the location in the orchard, and the location on the tree have a small effect on the flower microbiome.

## Materials and methods

### Collection site and weather

Samples were collected from twelve pear trees (*Pyrus communis* cultivars Spadona and Coscia) at an orchard in the Hula Valley in northern Israel (33.1505N, 35.6261E) during March and April 2019. The trees were ten years old. The plot has a history of fire blight, and inoculation experiments with *E. amylovora* were performed in previous years. The plot was treated with pesticides using the conventional local protocol in previous years but not in 2019. The undergrowth was sprayed with herbicides before the flowering period. Weather data (**Fig. S1**) was collected by a meteorological station adjacent to the orchard. The station provides meteorological data to alert farmers of fire blight favorable conditions based on the FBCA system (33). *E. amylovora* infection alerts were reported by the FBCA system multiple times during the flowering period.

### Sample collection strategy

The sample collection strategy and the number of samples that were collected are summarized in **Fig. 1**. For each cultivar, two groups of three neighboring trees were sampled (a total of six trees for each cultivar) to evaluate the variation in microbiome composition resulting from location within the orchard. The groups were located ∼15 meters apart. To test the variation in microbiome composition resulting from height, we collected flowers at heights ∼1.5 and ∼2.5 meters (**Fig. 1A**). Spadona flowers and leaves were collected on three dates between March 24th and March 31st, and Coscia samples were collected on five dates between April 11th and April 21st. All samples were collected during the morning hours. For the Coscia cultivar, flowers were sampled from 2-4 prevalent phenological stages in each collection date. For the Spadona cultivar, flowers of a single phenological stage were collected on each date (**Fig. 1B** and C). Overall, twelve samples were collected for each phenological stage at each date. One leaf sample was collected from each tree at all collection dates (six per collection date).

### Sample collection

30 flowers were collected for each flower sample, following (8), and 15 leaves were collected for each leaf sample. All flowers and leaves were collected using sterilized tweezers, and samples were immediately placed in liquid nitrogen, transported to the laboratory, and stored in -80°C until processing.

### DNA extraction and amplification

We applied the DNA extraction protocol described in (8) with minor modifications. The samples were processed a few weeks after collection. The extraction process included steps to separate the microbial cells from the plant material and to reduce the amount of chloroplasts and mitochondria. Samples were taken from the freezer and thawed on ice for ∼30 minutes. Next, we added 35ml 1x PBS-0.15% Tween solution, with 5-7 glass beads as a mild abrasive. The samples were then put into an orbital shaker at 270 rpm for 20 minutes at 4°C, followed by a 5-minute sonication in a water bath (Elmasonics S30H sonicator). The flower debris was removed using a sterile gause cloth into a new, sterile tube. The samples were centrifuged at 272g for 5 minutes (Eppendorf 5810R) at 4°C, to remove all remaining flower debris. The top liquid was centrifuged at 5111g for 30 minutes at 4°C. The pellet was collected, and the DNA was extracted using Exgene Cell DNA Isolation kit sv100 prep, following the manufacturer’s instructions.

Sequence libraries were prepared and sequenced at the Research Resources Center of the University of Illinois in Chicago. Libraries were prepared using the Nextera XT DNA Library Preparation kit and sequenced on Illumina MiSeq sequencer (2×250bp). The v3-v4 region of the 16S ribosomal small subunit was amplified using the 341f/785r primer set 341f (5’-CCTACGGGNGGCWGCAG-3’), 785r (5’-GACTACHVGGGTATCTAATC-3’). These primers were previously found to recover a high diversity of plant-associated bacteria (34). Samples with low sequence counts or high chloroplast contamination were dropped from the analysis. Overall, 176 flower samples (34 Spadona, 142 Coscia) and 47 leaf samples (17 Spadona, 30 Coscia) were analyzed (**Fig 1B**, **Table S1)**.

### Sequence analysis and statistics

All the initial analyses were performed using qiime2 v2022.8 (35). We processed the paired-end reads using the Dada2 plugin using the ‘pseudo’ option (36). 18,507 unique amplicon sequence variants (ASVs, also termed features) were extracted from 7,500,628 sequences that passed quality control, an average of 33,635 reads per sample.

Amplicon sequence variants were aligned using mafft (37), and phylogeny was constructed using fasttree (38). The taxonomy was assigned using the q2-feature-classifier plugin with the classify-sklearn option (39) against release 138 of the SILVA database (40), SSURef NR99 collection. Mitochondria and chloroplast sequences were filtered out. Some samples contained ASVs assigned to the Buchnera genus with 100% identity to *B. aphidicola*, an endosymbiont of aphids. These ASVs were removed from subsequent analysis. We also removed ASVs present at less than 0.0001% of the reads. Overall, 18,451 ASVs retained. The samples were rarefied (41) separately for each analysis using phyloseq. Refer to **Table S2** for exact numbers.

The downstream analyses, graphs, and statistics were done in R, using the phyloseq (version 1.36.0) (42) and the vegan (43) packages. Alpha diversity was computed through the Shannon index using phyloseq. Beta diversity was calculated using the vegan package. Constrained ordination was calculated with the ordinate function in phyloseq, using “CAP” method and Bray-Curtis distances. PERMANOVA (permutational multivariate analysis of variance) analysis (44) was computed using the adonis function in vegan (version 2.5.7) with 9,999 permutations. Comparisons of alpha diversities were done using the Kruskal-Wallis tests. To identify their origin, all *Erwiniaceae* ASVs were aligned against the NCBI nt database using BLAST (45). Differential abundance calculations were done using the corncob package in R (46).

## Acknowledgements

This work was supported by a grant from the JCA Foundation (Accelerator project). We thank Sondra Turjeman for her help with the statistical analyses. The authors acknowledge the assistance of the Northern Agriculture R&D experimental farm staff in the pear orchard maintenance.

## Code and data availability

Raw sequencing data for the project is available via SRA with BioProject identifier PRJNA1043619. The code used for the analyses in this paper is available from https://github.com/SharonLab/pear-microbiome.

## References

1. Trivedi P, Leach JE, Tringe SG, Sa T, Singh BK. Plant-microbiome interactions: from community assembly to plant health. Nat Rev Microbiol. 2020 Nov;18(11):607–21.

2. Brader G, Compant S, Vescio K, Mitter B, Trognitz F, Ma L-J, et al. Ecology and Genomic Insights into Plant-Pathogenic and Plant-Nonpathogenic Endophytes. Annu Rev Phytopathol. 2017 Aug 4;55:61–83.

3. Compant S, Samad A, Faist H, Sessitsch A. A review on the plant microbiome: Ecology, functions, and emerging trends in microbial application. J Advanc Res. 2019 Sep;19:29– 37.

4. Liu H, Macdonald CA, Cook J, Anderson IC, Singh BK. An Ecological Loop: Host Microbiomes across Multitrophic Interactions. Trends Ecol Evol. 2019 Dec;34(12):1118– 30.

5. Rastogi G, Coaker GL, Leveau JHJ. New insights into the structure and function of phyllosphere microbiota through high-throughput molecular approaches. FEMS Microbiol Lett. 2013 Nov;348(1):1–10.

6. Mansfield J, Genin S, Magori S, Citovsky V, Sriariyanum M, Ronald P, et al. Top 10 plant pathogenic bacteria in molecular plant pathology. Mol Plant Pathol. 2012 Aug;13(6):614– 29.

7. Vega C, Álvarez-Pérez S, Albaladejo RG, Steenhuisen S, Lachance M, Johnson SD, et al. The role of plant–pollinator interactions in structuring nectar microbial communities. J Ecol. 2021 Sep;109(9):3379–95.

8. Shade A, McManus PS, Handelsman J. Unexpected diversity during community succession in the apple flower microbiome. MBio. 2013 Feb 26;4(2).

9. Cui Z, Huntley RB, Schultes NP, Steven B, Zeng Q. Inoculation of Stigma-Colonizing Microbes to Apple Stigmas Alters Microbiome Structure and Reduces the Occurrence of Fire Blight Disease. Phytobiomes Journal. 2021 Jan 14;PBIOMES-04-20-0.

10. Vannette RL, Fukami T. Contrasting effects of yeasts and bacteria on floral nectar traits. Ann Bot. 2018 Jun 8;121(7):1343–9.

11. Helletsgruber C, Dötterl S, Ruprecht U, Junker RR. Epiphytic bacteria alter floral scent emissions. J Chem Ecol. 2017 Dec;43(11–12):1073–7.

12. Liu H, Brettell LE, Singh B. Linking the phyllosphere microbiome to plant health. Trends Plant Sci. 2020 Sep;25(9):841–4.

13. Smessaert J, Van Geel M, Verreth C, Crauwels S, Honnay O, Keulemans W, et al. Temporal and spatial variation in bacterial communities of “Jonagold” apple (Malus x domestica Borkh.) and “Conference” pear (Pyrus communis L.) floral nectar. Microbiologyopen. 2019 Dec;8(12):e918.

14. Vanneste JL. What is fire blight? Who is Erwinia amylovora? How to control it? In: Vanneste JL, editor. Fire blight: the disease and its causative agent, Erwinia amylovora. Wallingford: CABI; 2000. p. 1–6.

15. Vanneste JL, Eden-Green S. Migration of Erwinia amylovora in host plant tissues. In: Vanneste JL, editor. Fire blight: the disease and its causative agent, Erwinia amylovora. Wallingford: CABI; 2000. p. 73–83.

16. van der Zwet T. Present worldwide distribution of fire blight. Acta Hortic. 1996 Apr;(411):7– 8.

17. Shtienberg D, Manulis-Sasson S, Zilberstaine M, Oppenheim D, Shwartz H. The incessant battle against fire blight in pears: 30 years of challenges and successes in managing the disease in israel. Plant Dis. 2015 Aug;99(8):1048–58.

18. Billing E. Fire blight. Why do views on host invasion by Erwinia amylovora differ? Plant Pathol. 2011 Apr;60(2):178–89.

19. Dafny-Yelin M, Moy JC, Stern RA, Doron I, Silberstein M, Michaeli D. High-density ‘Spadona’ pear orchard shows reduced tree sensitivity to fire blight damage due to decreased tree vigour. Phytopathol Mediterr. 2021 Nov 15;60(3):421–6.

20. Billing E. Fire blight risk assessment: billing’s integrated system (bis) and its evaluation. Acta Hortic. 1999 Jul;(489):399–406.

21. Farkas Á, Mihalik E, Dorgai L, Bubán T. Floral traits affecting fire blight infection and management. Trees. 2012 Feb;26(1):47–66.

22. Billing E. Billing’s revised system (BRS) for fireblight risk assessment. EPPO Bulletin. 1992 Mar;22(1):1–102.

23. Shtienberg D, Shwartz H, Oppenheim D, Zilberstaine M, Herzog Z, Manulis S, et al. Evaluation of local and imported fire blight warning systems in Israel. Phytopathology. 2003 Mar;93(3):356–63.

24. Cui Z, Huntley RB, Zeng Q, Steven B. Temporal and spatial dynamics in the apple flower microbiome in the presence of the phytopathogen Erwinia amylovora. ISME J. 2021 Jan;15(1):318–29.

25. Russo NL, Burr TJ, Breth DI, Aldwinckle HS. Isolation of Streptomycin-Resistant Isolates of Erwinia amylovora in New York. Plant Dis. 2008 May;92(5):714–8.

26. Dafny-Yelin M, Moy JC, Mairesse O, Silberstein M, Sapir G, Michaeli D. Efficacy of fire blight management in pome fruit in northern Israel: copper agents and their effect on yield parameters. J Plant Pathol. 2021 Aug;103(S1):151–61.

27. Schaeffer RN, Pfeiffer VW, Basu S, Brousil M, Strohm C, DuPont ST, et al. Orchard management and landscape context mediate the pear floral microbiome. Appl Environ Microbiol. 2021 Jul 13;87(15):e0004821.

28. Ait Bahadou S, Ouijja A, Karfach A, Tahiri A, Lahlali R. New potential bacterial antagonists for the biocontrol of fire blight disease (Erwinia amylovora) in Morocco. Microb Pathog. 2018 Apr;117:7–15.

29. Steven B, Huntley RB, Zeng Q. The influence of flower anatomy and apple cultivar on the apple flower phytobiome. Phytobiomes. 2018 Jul 25;PBIOMES-03-18-0.

30. Kim WS, Gardan L, Rhim SL, Geider K. Erwinia pyrifoliae sp. nov., a novel pathogen that affects Asian pear trees (Pyrus pyrifolia Nakai). Int J Syst Bacteriol. 1999 Apr;49 Pt 2:899– 905.

31. Gaube P, Junker RR, Keller A. Changes amid constancy: flower and leaf microbiomes along land use gradients and between bioregions. BioRxiv. 2020 Apr 1;

32. Santander RD, Biosca EG. Erwinia amylovora psychrotrophic adaptations: evidence of pathogenic potential and survival at temperate and low environmental temperatures. PeerJ. 2017 Oct 26;5:e3931.

33. Shtienberg D, Kritzman G, Herzog Z, Openhaim D, Zillberstein M, Blatchinsky D. Development and evaluation of a decision support system for management of fire blight in pears. Acta Hortic. 1999 Jul;(489):385–92.

34. Thijs S, Op De Beeck M, Beckers B, Truyens S, Stevens V, Van Hamme JD, et al. Comparative Evaluation of Four Bacteria-Specific Primer Pairs for 16S rRNA Gene Surveys. Front Microbiol. 2017 Mar 28;8:494.

35. Bolyen E, Rideout JR, Dillon MR, Bokulich NA, Abnet CC, Al-Ghalith GA, et al. Reproducible, interactive, scalable and extensible microbiome data science using QIIME 2. Nat Biotechnol. 2019 Aug;37(8):852–7.

36. Callahan BJ, McMurdie PJ, Rosen MJ, Han AW, Johnson AJA, Holmes SP. DADA2: High-resolution sample inference from Illumina amplicon data. Nat Methods. 2016 May 23;13(7):581–3.

37. Katoh K, Misawa K, Kuma K, Miyata T. MAFFT: a novel method for rapid multiple sequence alignment based on fast Fourier transform. Nucleic Acids Res. 2002 Jul 15;30(14):3059–66.

38. Price MN, Dehal PS, Arkin AP. FastTree 2 — approximately maximum-likelihood trees for large alignments. PLoS ONE. 2010 Mar 10;5(3):e9490.

39. Bokulich NA, Kaehler BD, Rideout JR, Dillon M, Bolyen E, Knight R, et al. Optimizing taxonomic classification of marker-gene amplicon sequences with QIIME 2’s q2-feature-classifier plugin. Microbiome. 2018 May 17;6(1):90.

40. Quast C, Pruesse E, Yilmaz P, Gerken J, Schweer T, Yarza P, et al. The SILVA ribosomal RNA gene database project: improved data processing and web-based tools. Nucleic Acids Res. 2013 Jan;41(Database issue):D590-6.

41. Weiss S, Xu ZZ, Peddada S, Amir A, Bittinger K, Gonzalez A, et al. Normalization and microbial differential abundance strategies depend upon data characteristics. Microbiome. 2017 Mar 3;5(1):27.

42. McMurdie PJ, Holmes S. phyloseq: an R package for reproducible interactive analysis and graphics of microbiome census data. PLoS ONE. 2013 Apr 22;8(4):e61217.

43. Oksanen J, Blanchet FG, Kindt R, Legendre P, Minchin PR, O’hara RB, et al. vegan: Community Ecology Package. R package version 2.2–1. 2015;

44. McArdle BH, Anderson MJ. Fitting multivariate models to community data: a comment on distance-based redundancy analysis. Ecology. 2001 Jan;82(1):290–7.

45. McGinnis S, Madden TL. BLAST: at the core of a powerful and diverse set of sequence analysis tools. Nucleic Acids Res. 2004 Jul 1;32(Web Server issue):W20-5.

46. Martin BD, Witten D, Willis AD. Modeling microbial abundances and dysbiosis with beta-binomial regression. Ann Appl Stat. 2020 Mar;14(1):94–115.

